## Supplementary figures for "Mechanisms underlying microglial colonization of developing neural retina in zebrafish"

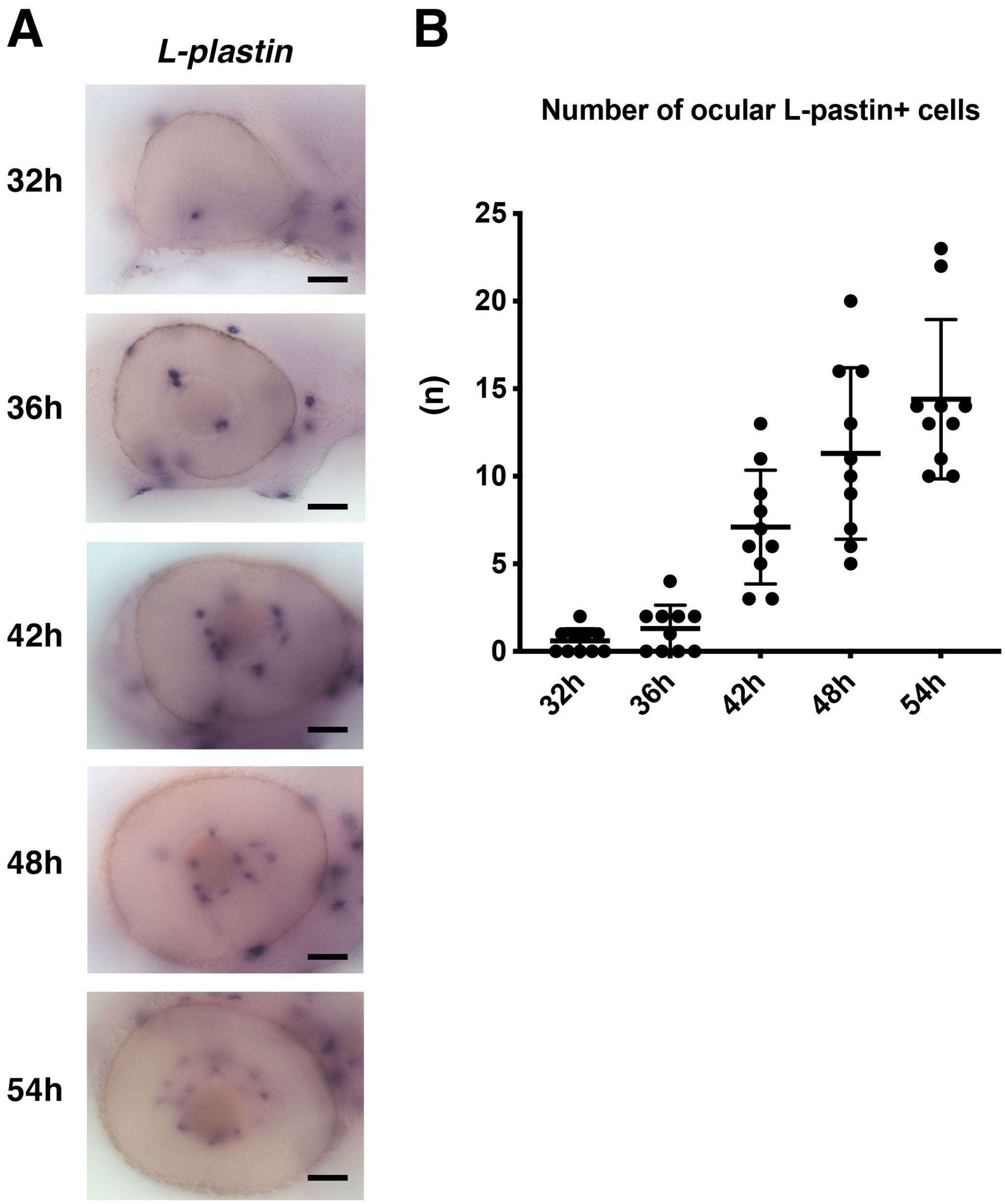


Fig. 1-figure supplement 1

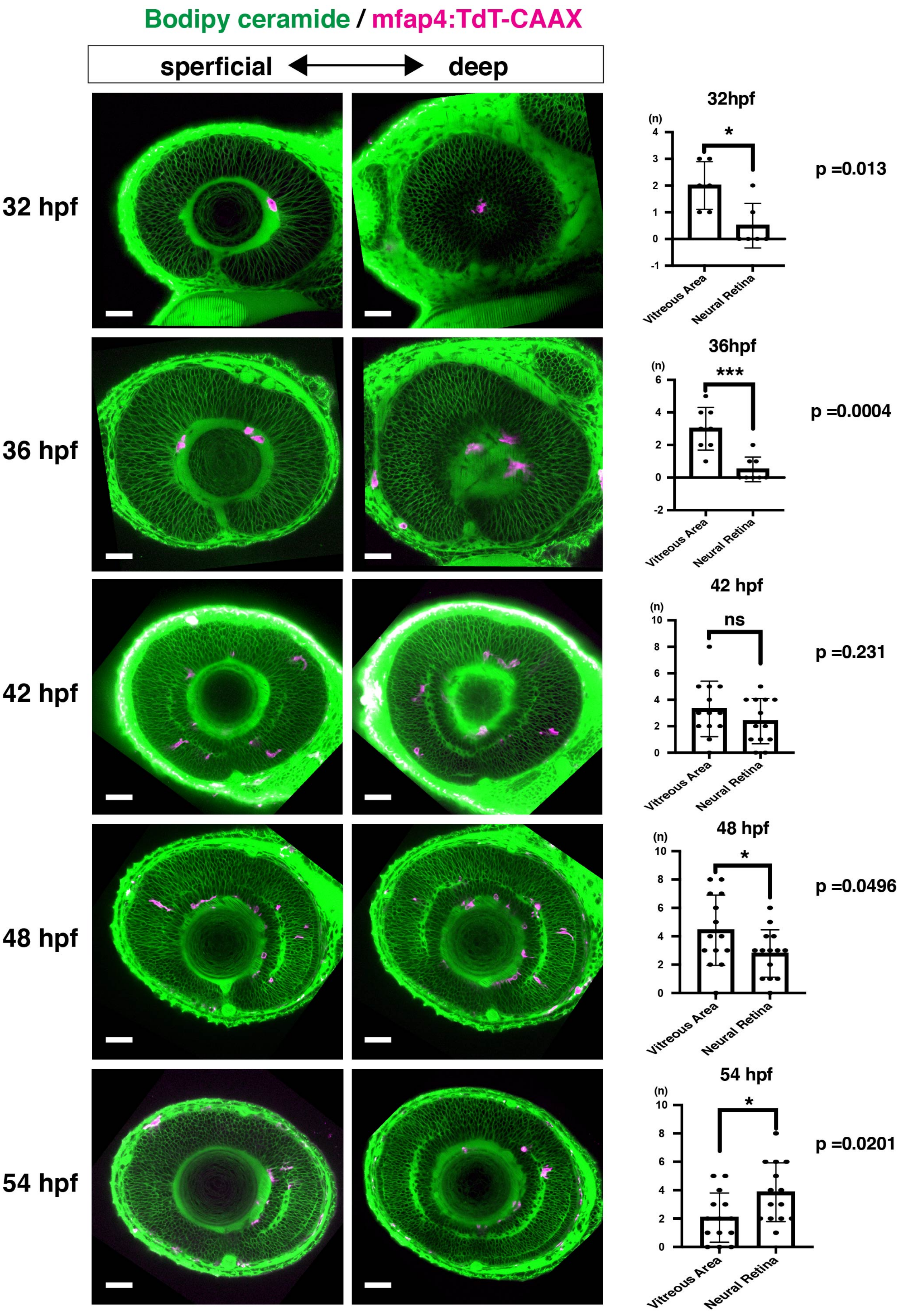


### Fig. 1-figure supplement 2

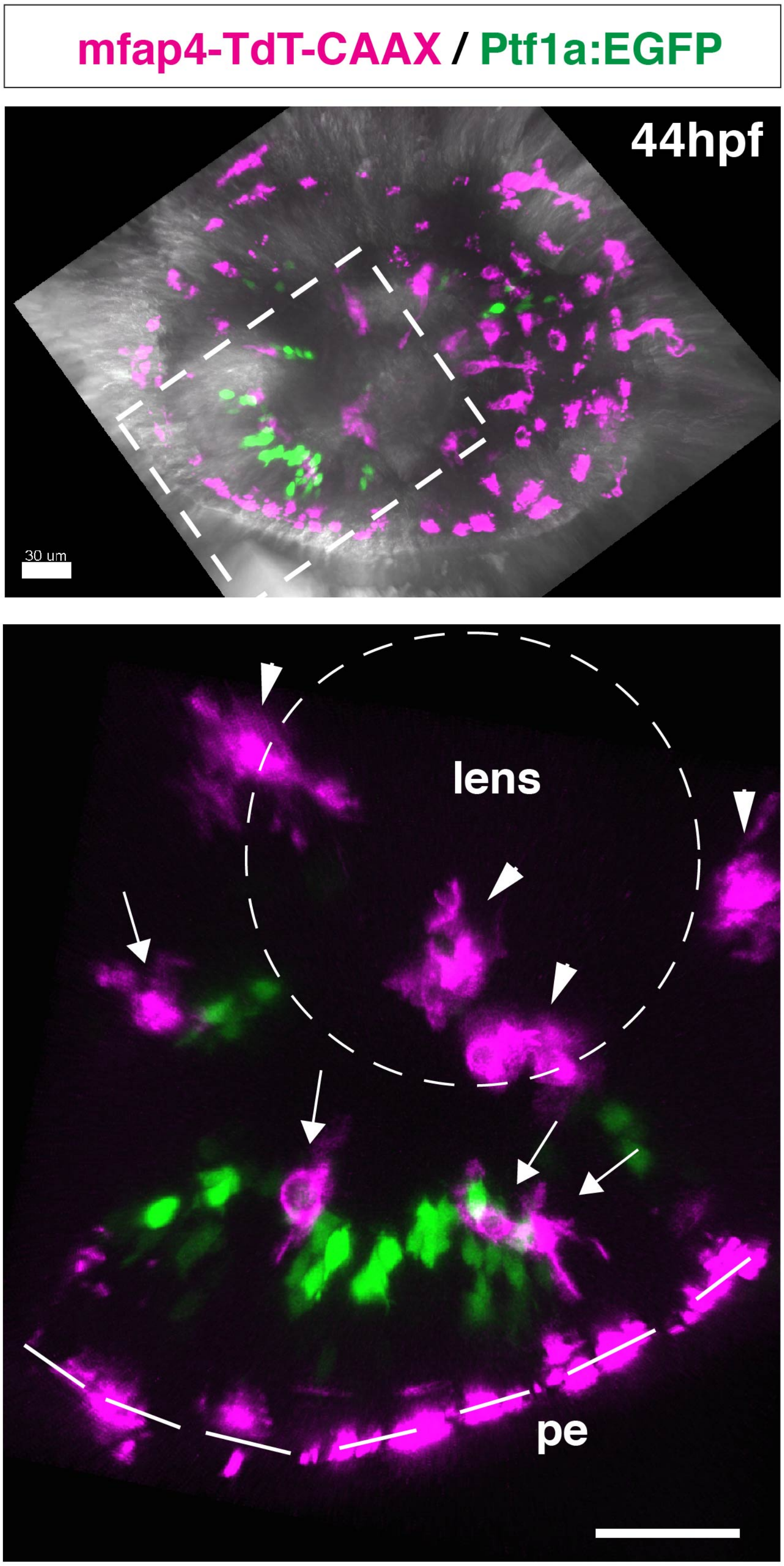


**Fig 1-figure supplement 3**

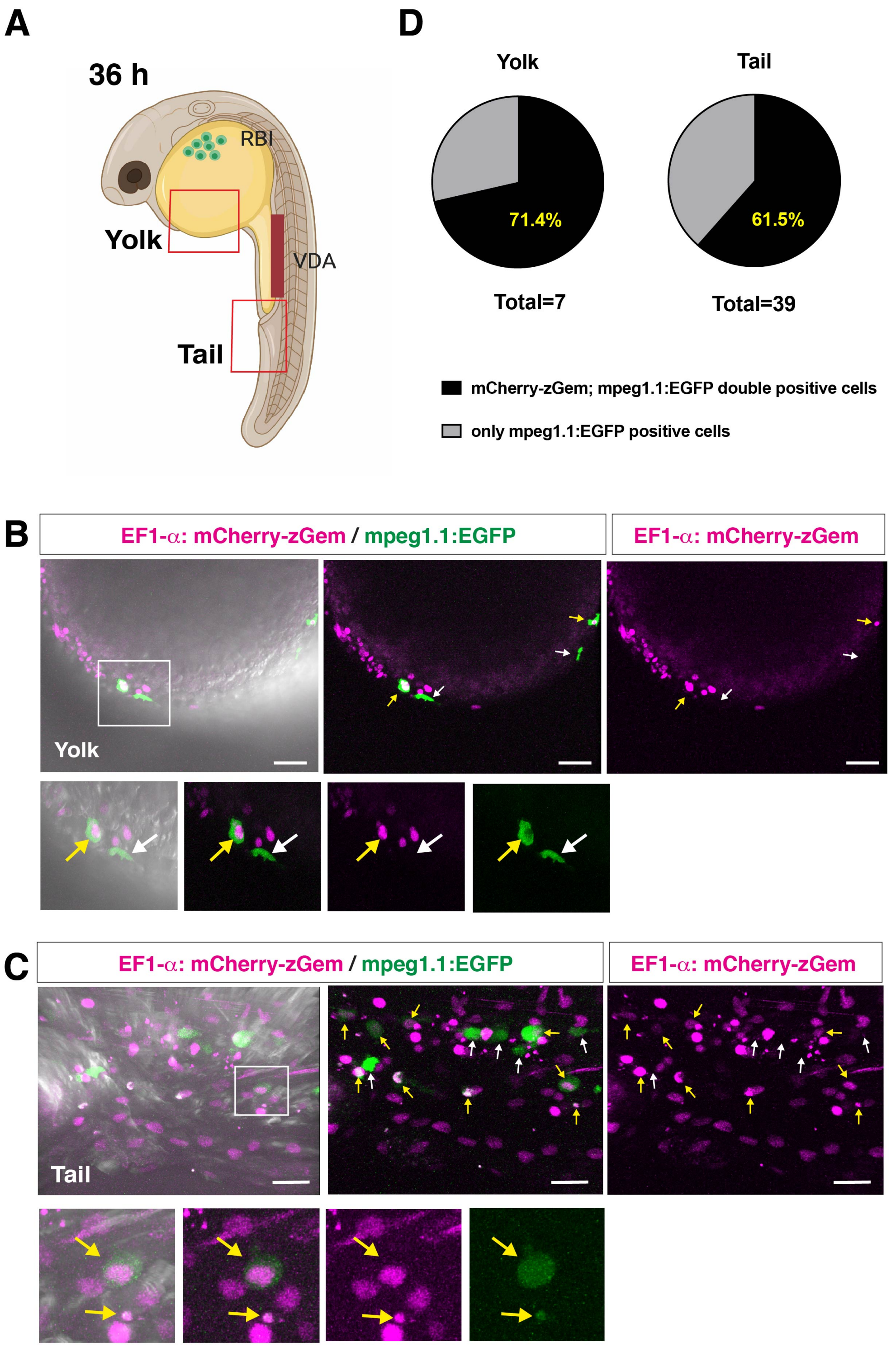


Fig.1 -figure supplement 4

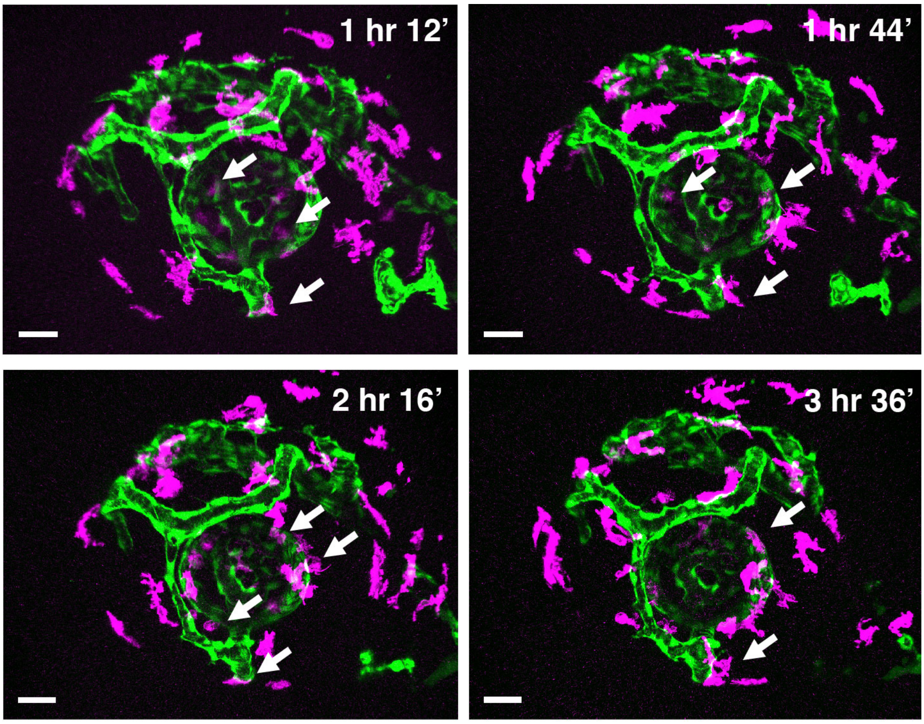


### Fig. 2-figure supplement 1

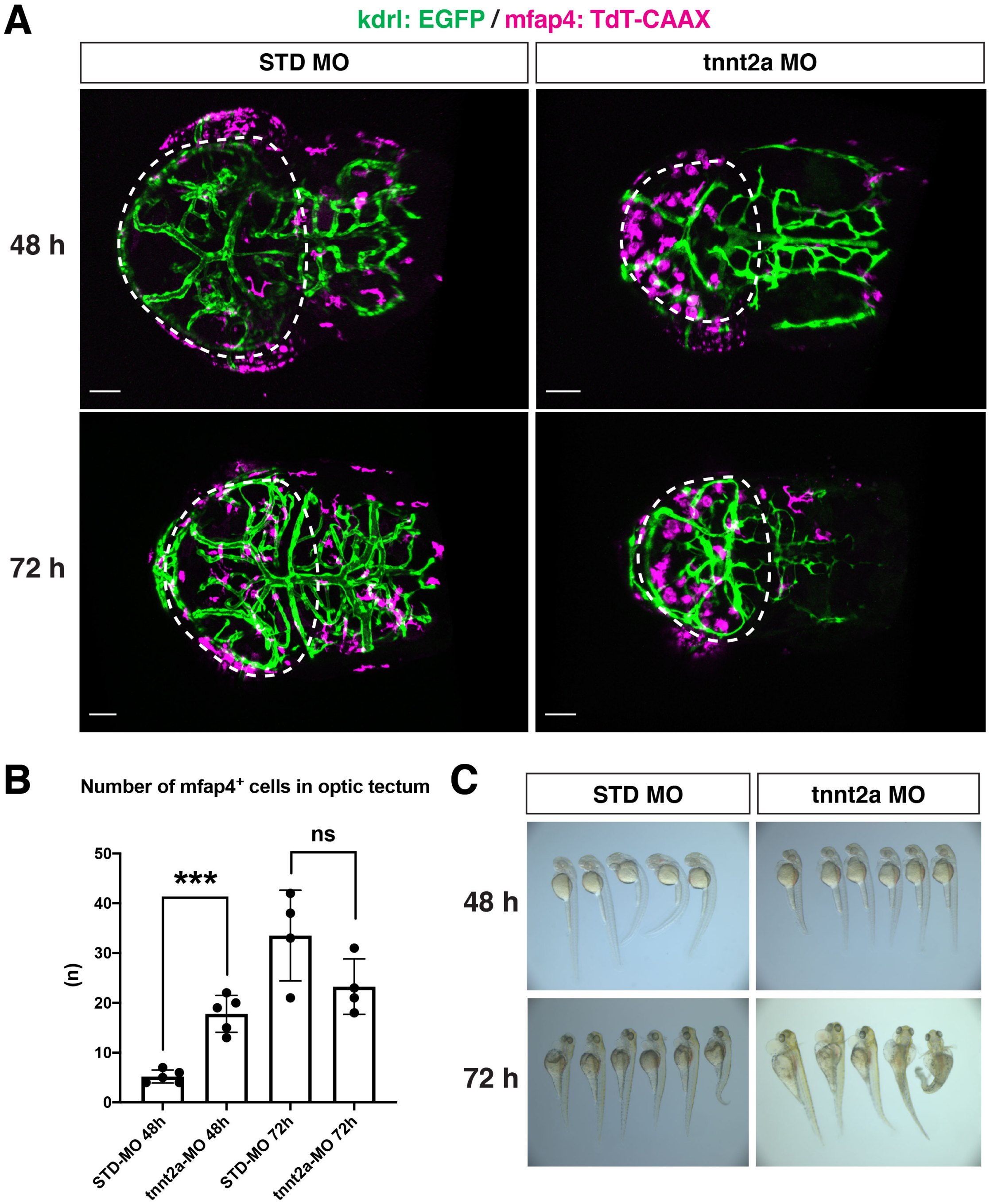


### Fig. 2-figure supplement 2

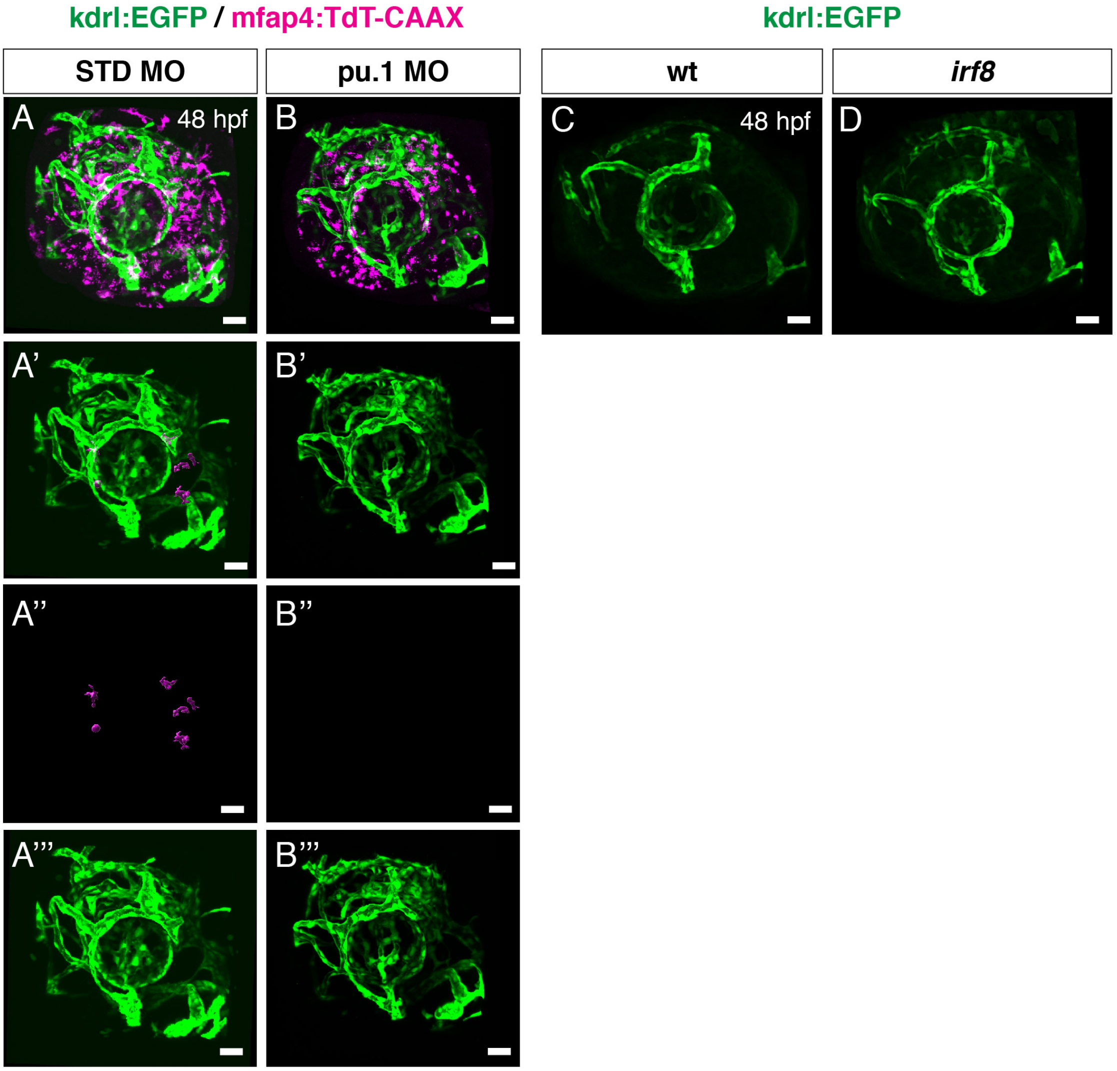


**Fig. 2-figure supplement 3**

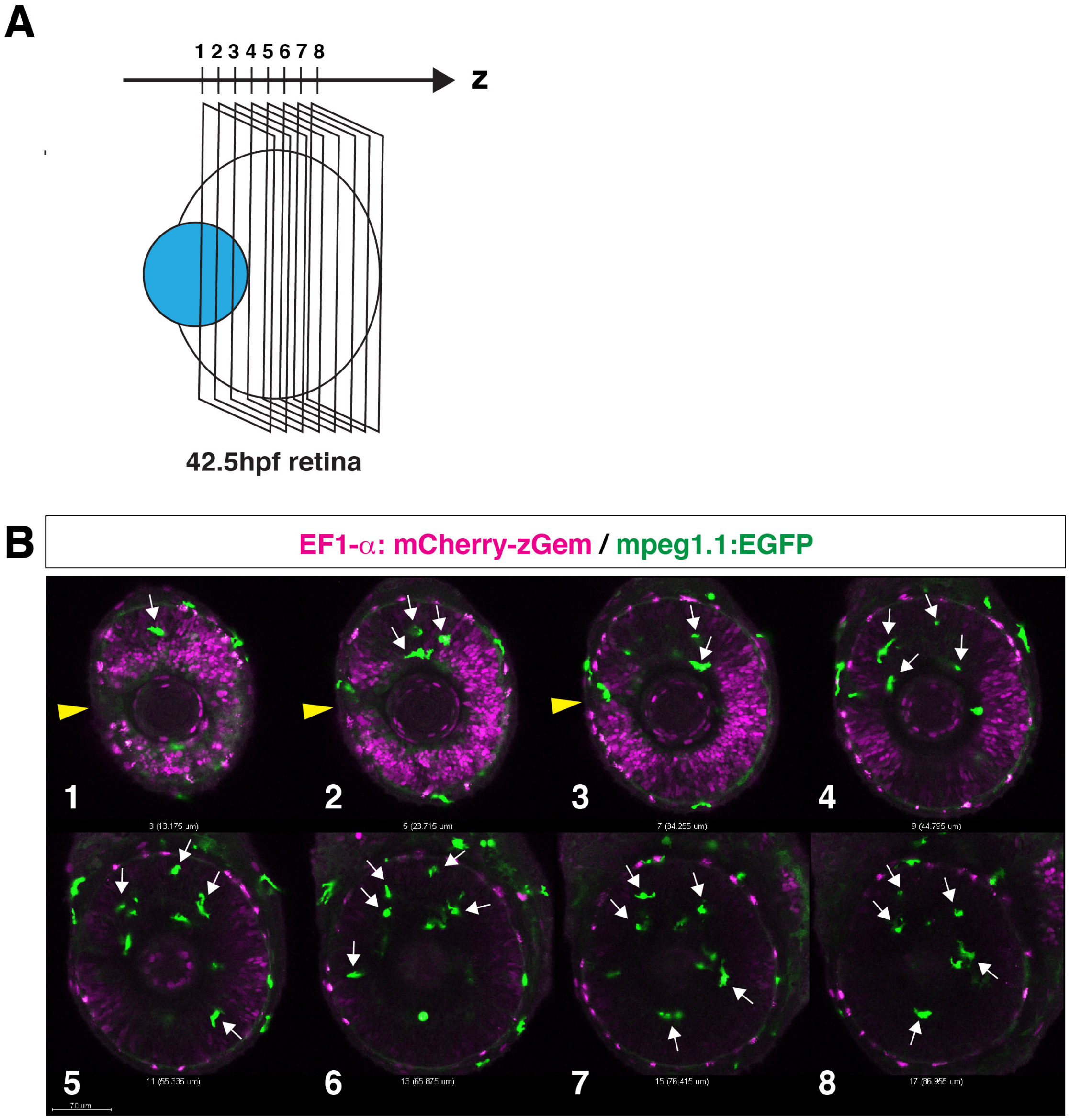


**Fig. 3-figure supplement 1**

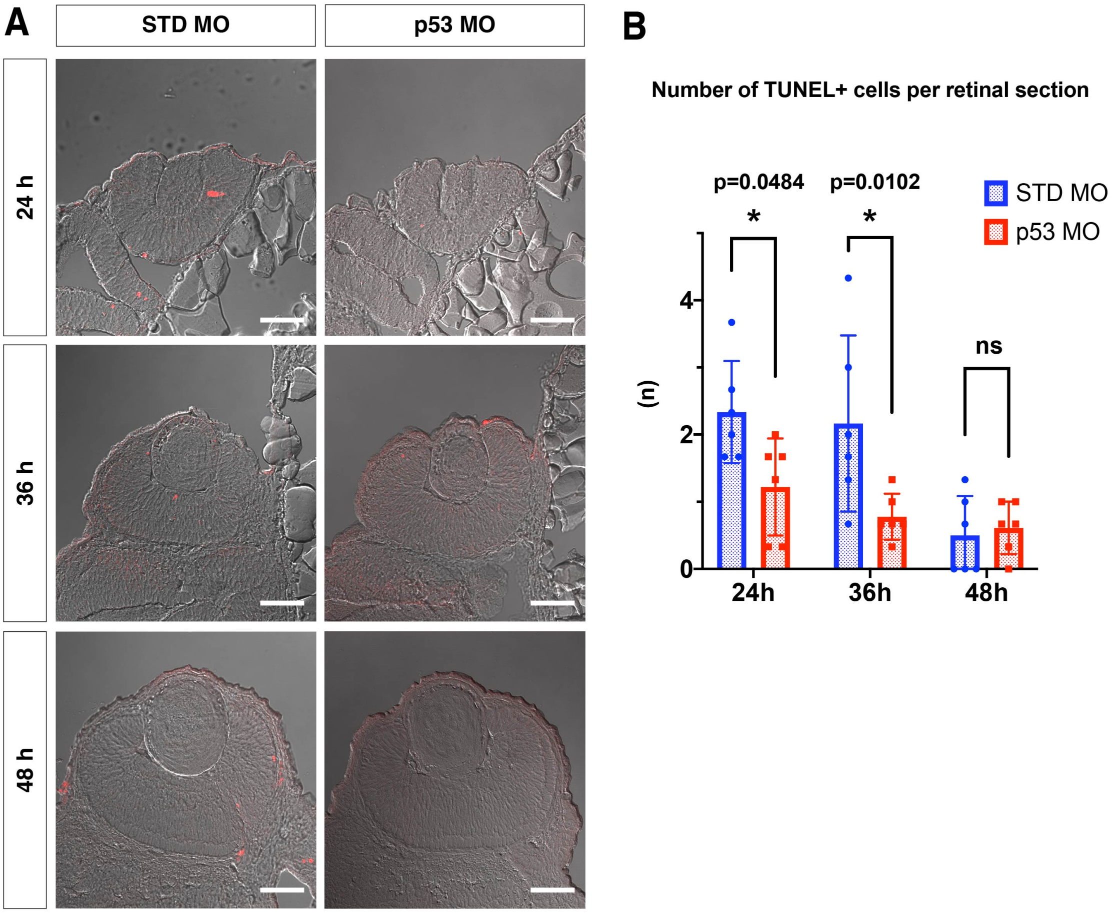


### Fig 3. Figure Supplement 2

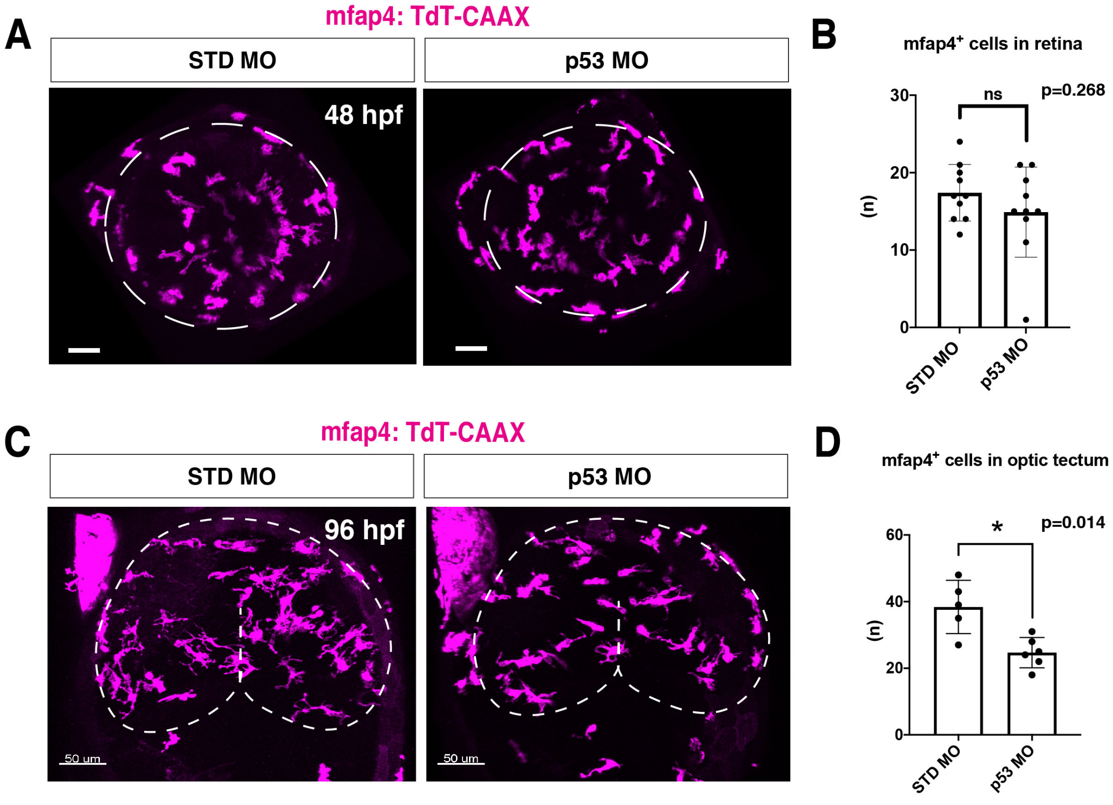


### Fig. 3-figure supplement 3

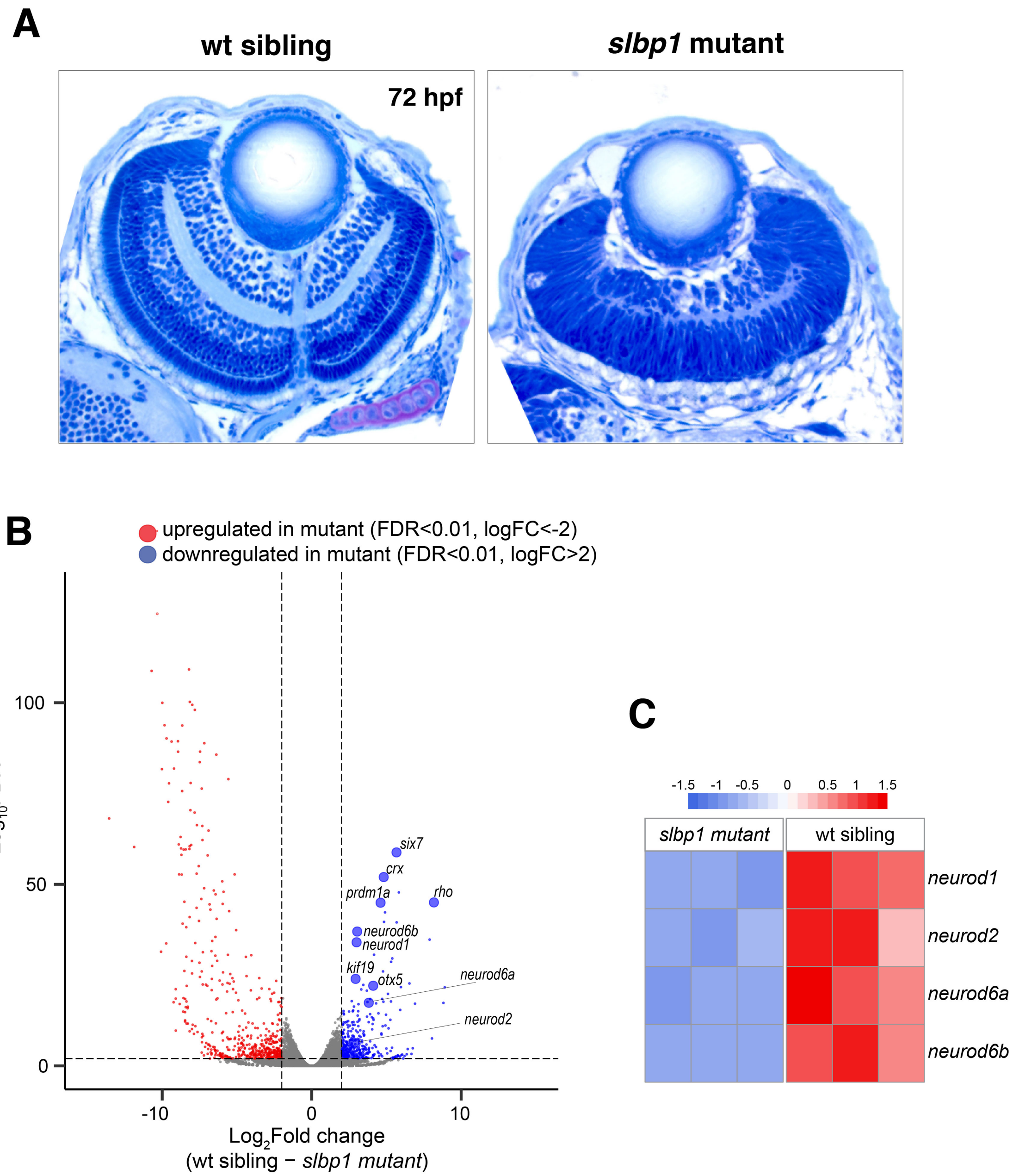


**Fig. 4-figure supplement 1**

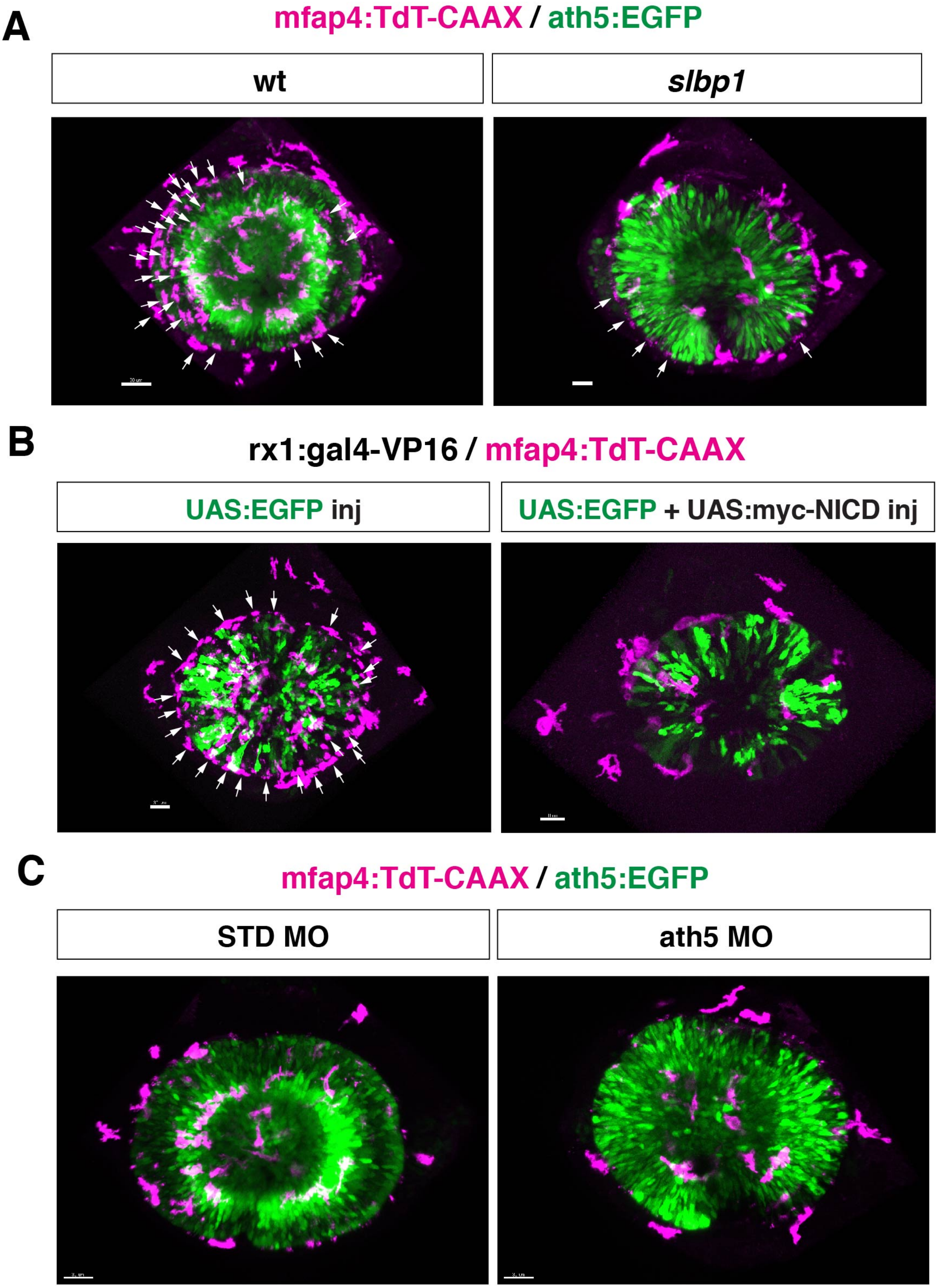


**Fig.4-figure supplement 2**

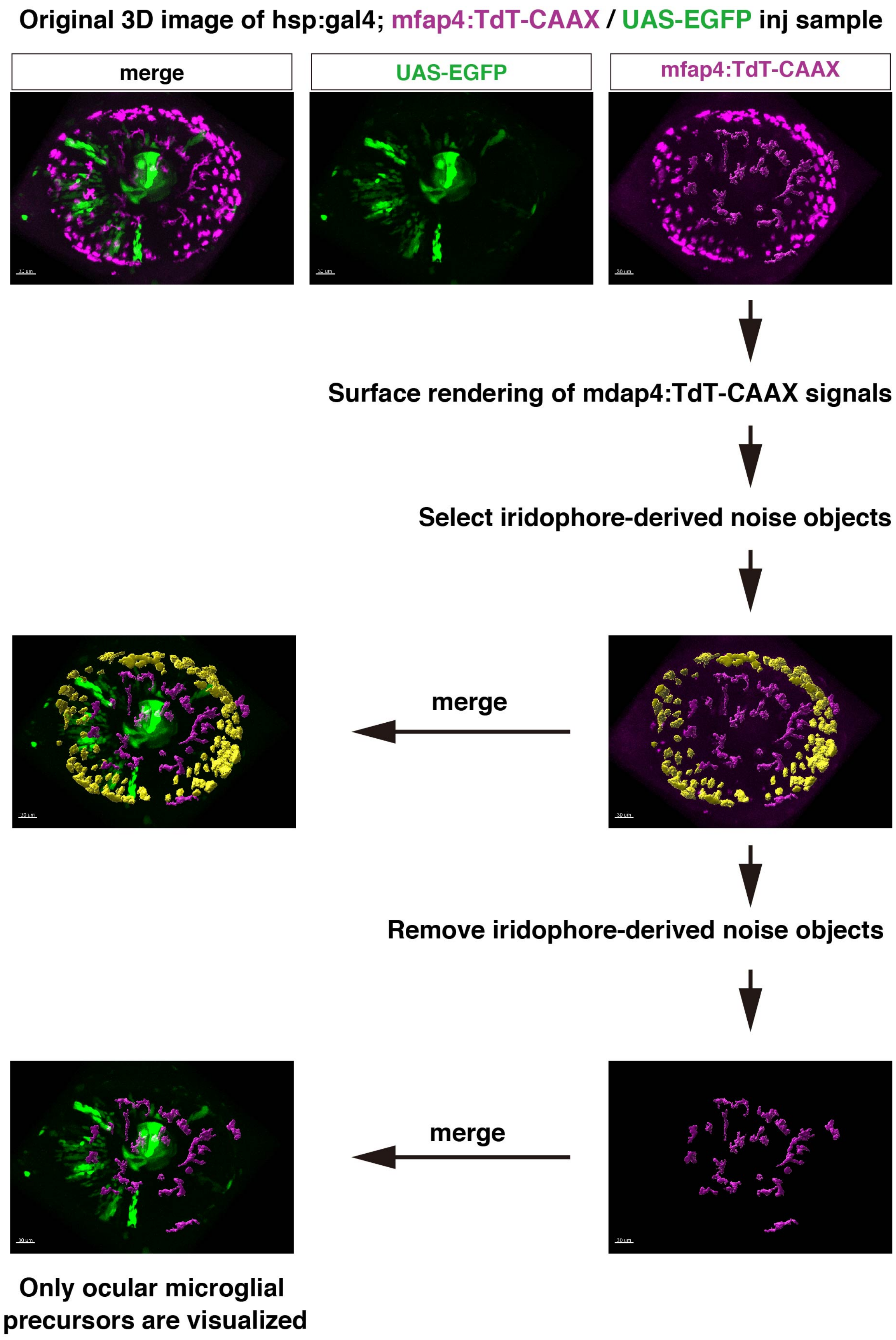


**Fig. 4-figure supplement 3**

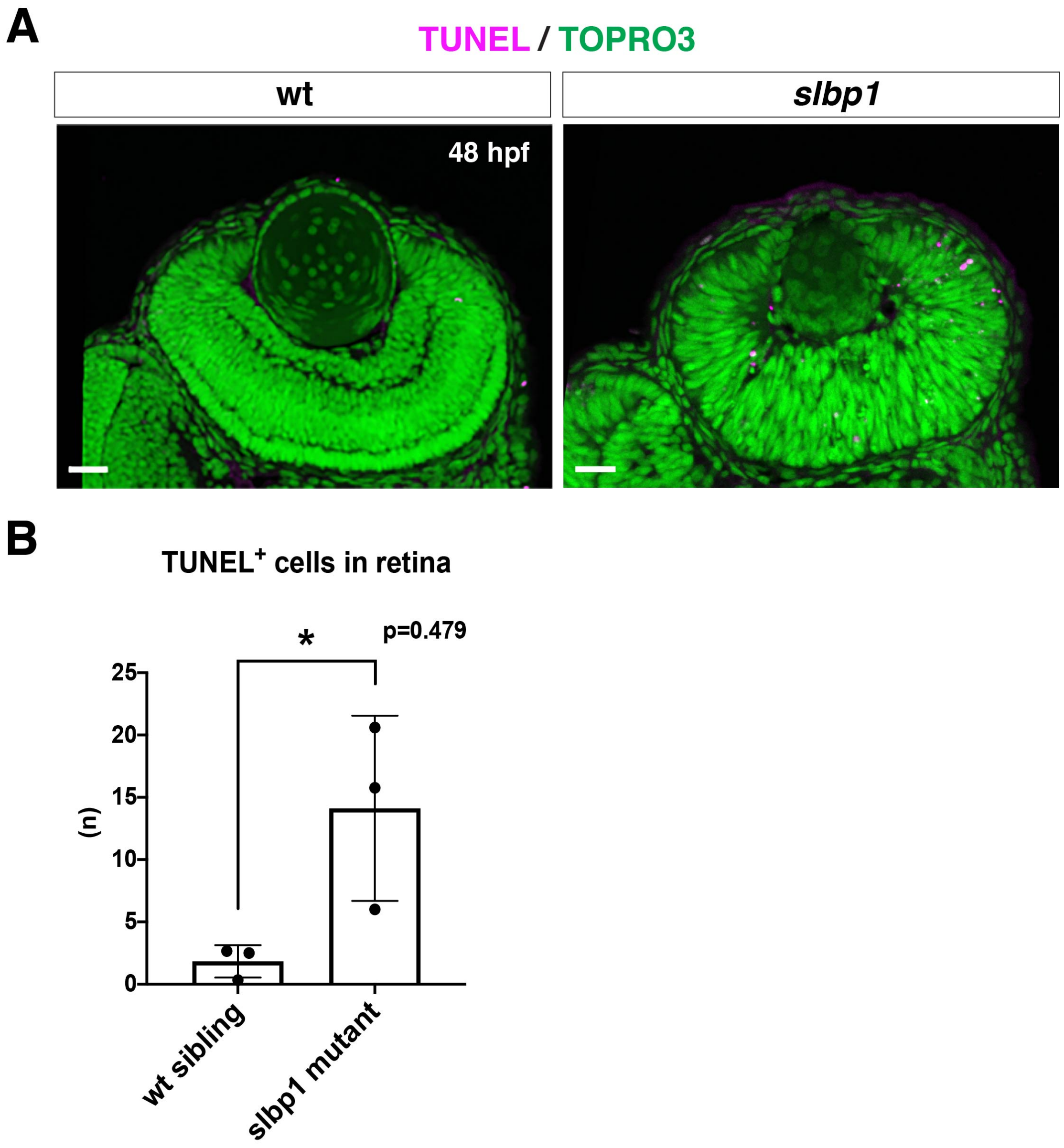


**Fig 4.-figure supplement 4**

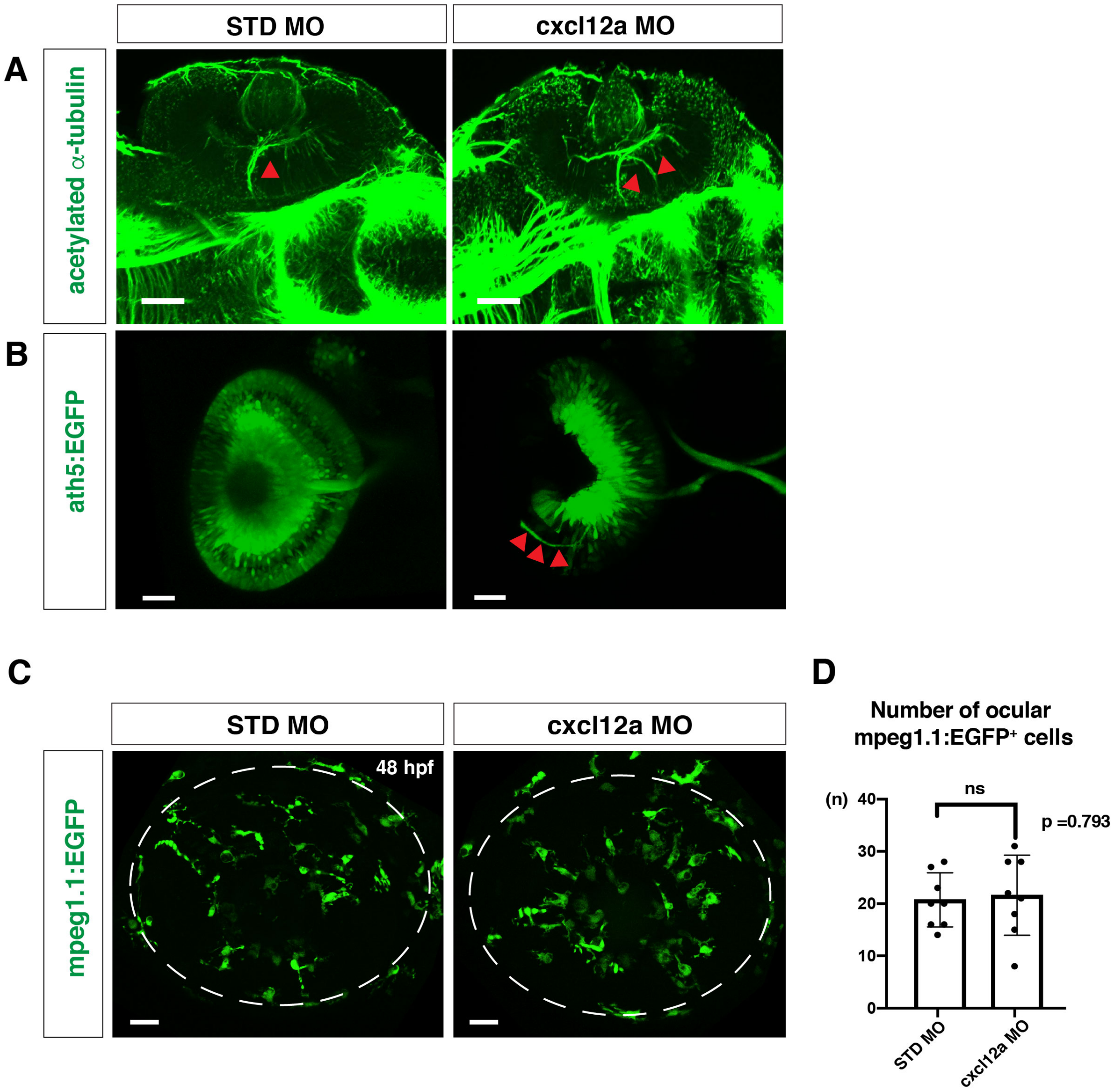


**Fig. 4-figure supplement 5**

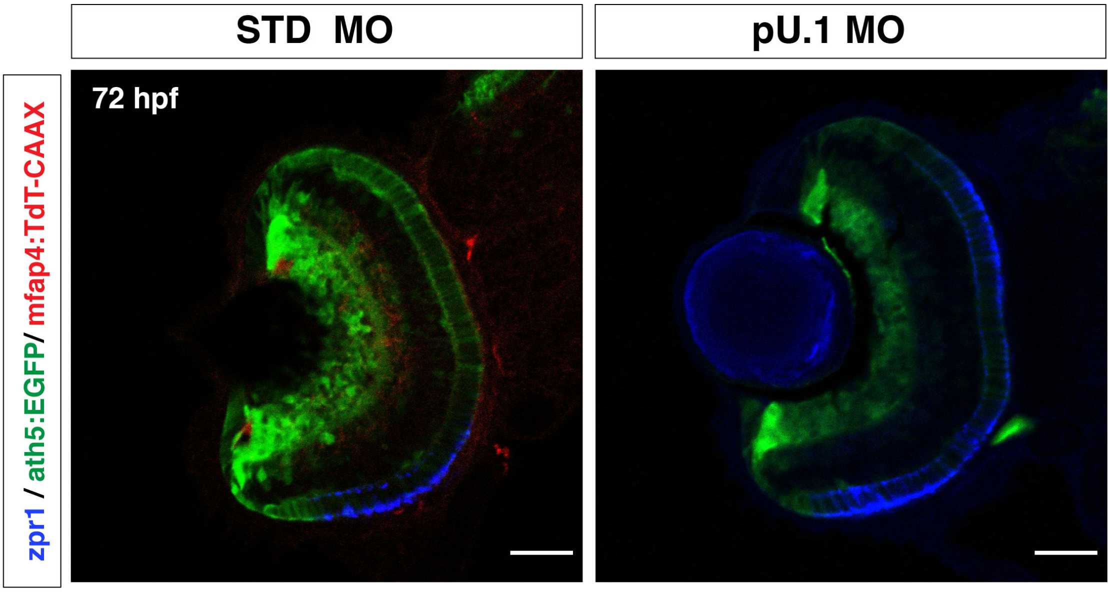


**Fig. 4-figure supplement 6**

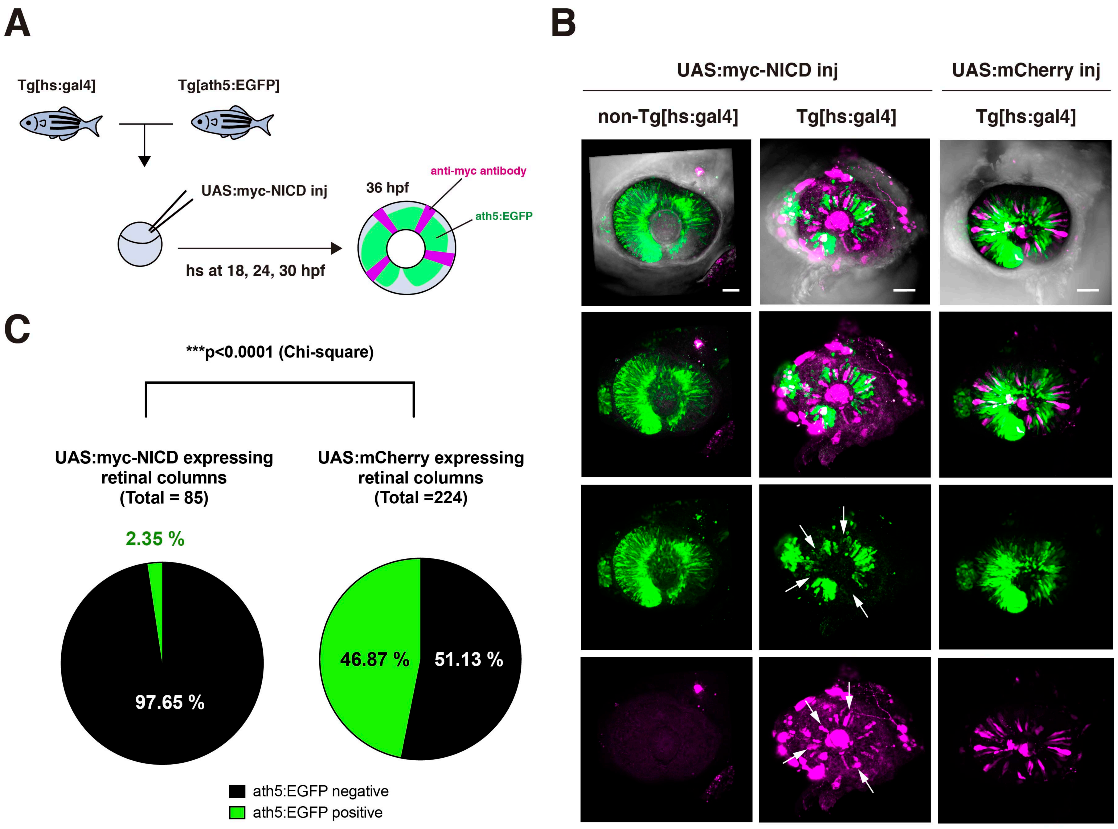


**Fig. 4- figure supplement 7**

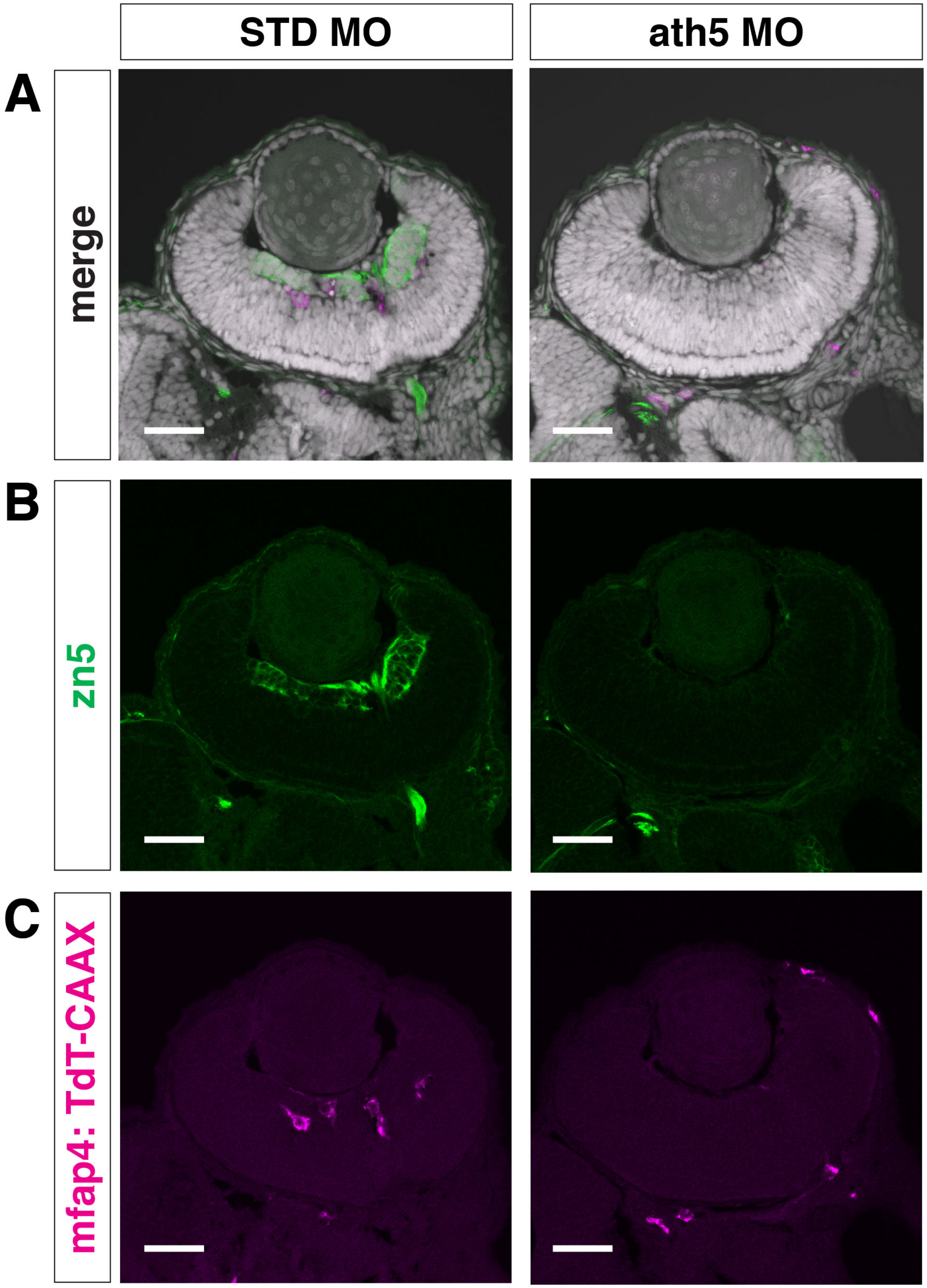


Fig. 4-figure supplement 8

A

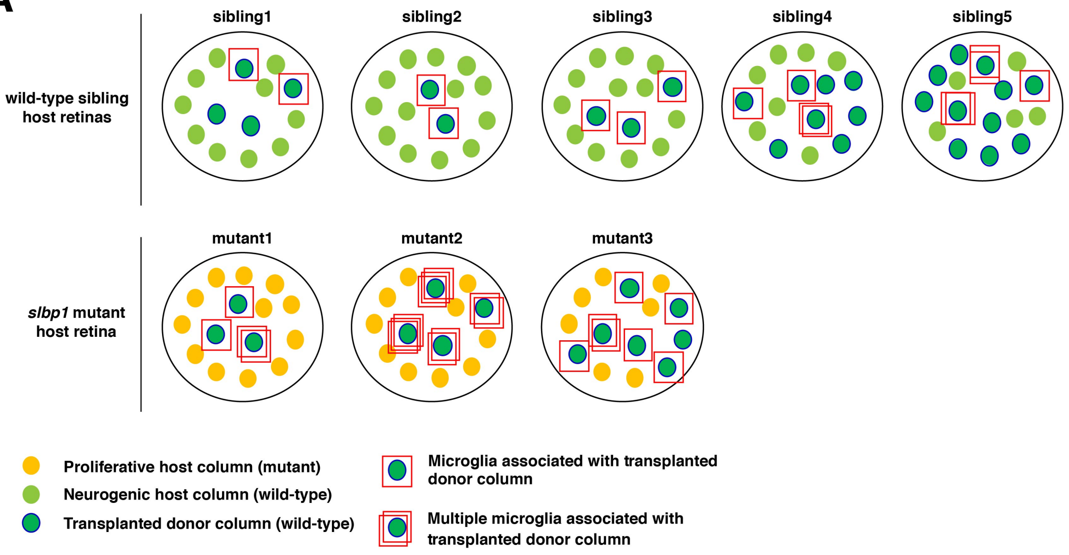


B

| Sample (host retina) | Total number of host microglial precursors | Number of host microglial precursors accosiated with transplanted donor columns | Fraction of host microglial precursors associated with transplanted donor columns (%) | Total number of transplanted donor columns | Trappiung efficiency of microglial precursors per transplaned donor colum (%) |
|---|---|---|---|---|---|
| sibling1 | 24 | 2 | 8.33 | 4 | 2.0825 |
| sibling2 | 22 | 2 | 9.091 | 2 | 4.5455 |
| sibling3 | 20 | 3 | 15 | 3 | 5 |
| sibling4 | 15 | 4 | 26.67 | 8 | 3.325 |
| sibling5 | 17 | 5 | 29.41 | 11 | 2.674 |
| mutant1 | 15 | 4 | 26.67 | 3 | 8.89 |
| mutant2 | 11 | 10 | 90.91 | 4 | 22.73 |
| mutant3 | 7 | 7 | 100 | 7 | 14.29 |

C

| Sample (injection construct) | Total number of ocular microglial precursors | Number of microglial precursors accosiated with EGFP-expressing columns | Fraction of microglial precursors associated with EGFP-expresssing columns (%) | Total number of EGFP-expressing columns | Trappiung efficiency of microglial precursors per EGFP-expressing colum (%) |
|---|---|---|---|---|---|
| UAS:EGFP inj 1 | 23 | 9 | 39 | 13 | 3 |
| UAS:EGFP inj 2 | 27 | 13 | 48 | 9 | 5.3 |
| UAS:EGFP inj 3 | 31 | 16 | 51 | 13 | 3.2 |
| UAS:EGFP inj 4 | 26 | 13 | 50 | 18 | 3.8 |
| UAS:EGFP inj 5 | 16 | 10 | 62.5 | 11 | 6.25 |
| UAS:EGFP+UAS:myc-NICD inj 1 | 14 | 3 | 21.4 | 7 | 3.06 |
| UAS:EGFP+UAS:myc-NICD inj 2 | 15 | 5 | 33.3 | 20 | 1.9 |
| UAS:EGFP+UAS:myc-NICD inj 3 | 8 | 2 | 25 | 23 | 1.08 |
| UAS:EGFP+UAS:myc-NICD inj 4 | 11 | 2 | 18.1 | 10 | 1.8 |
| UAS:EGFP+UAS:myc-NICD inj 5 | 18 | 3 | 16.6 | 26 | 1.29 |

**Fig. 5 figure supplement 1**

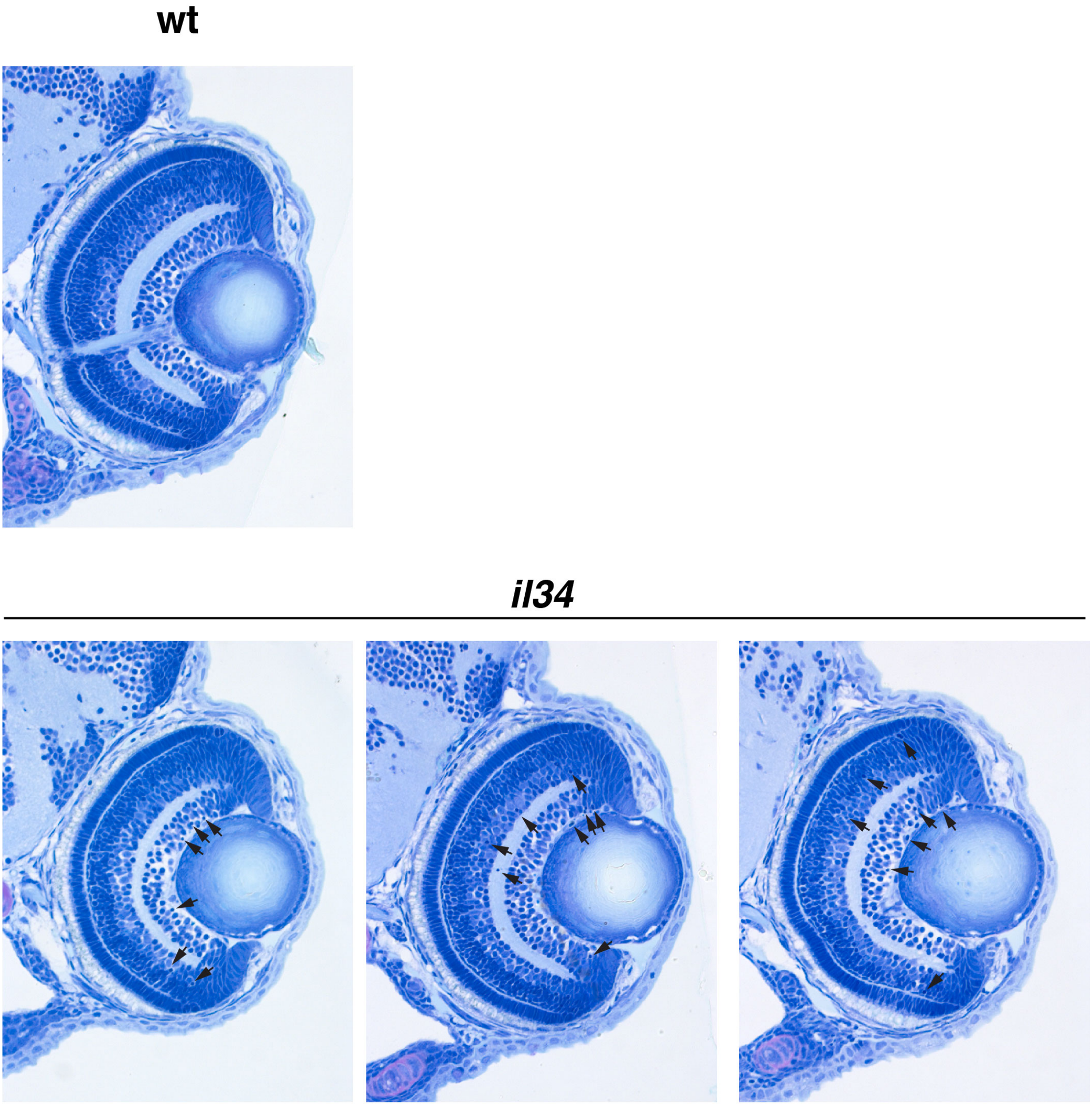


### Fig. 6-figure supplement 1

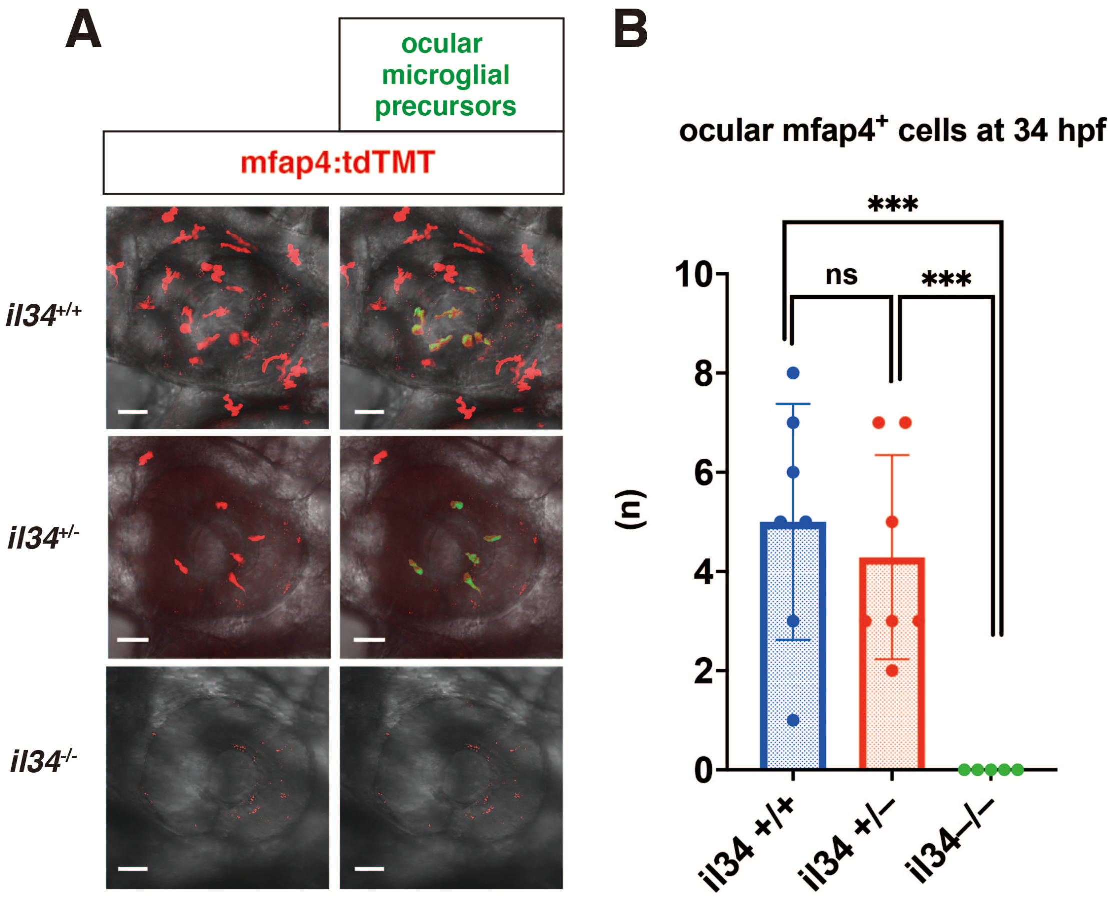


**Fig. 6-figure supplement 2**

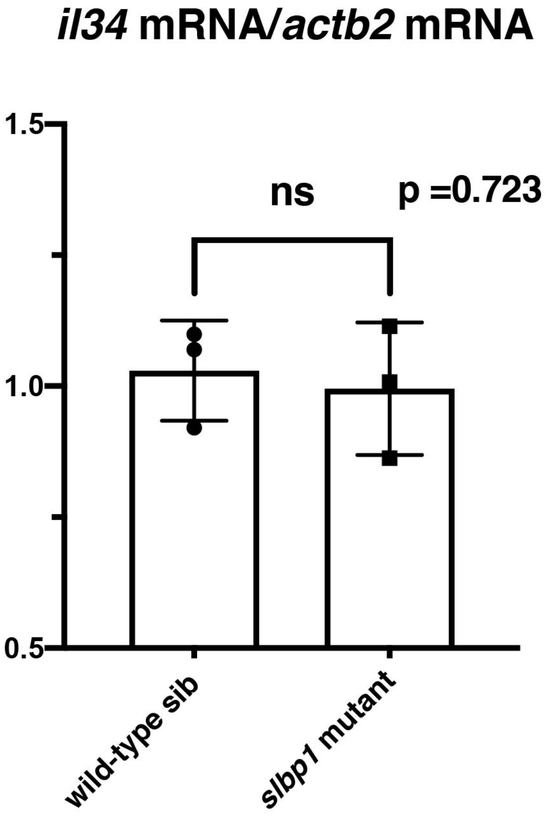


**Fig. 6-figure supplement 3**
